# Cells stably expressing shRNA against MYO10 display altered cell motility

**DOI:** 10.1101/2025.07.23.666455

**Authors:** Joanna A. Mas, Chase E. Cristella, Vu M.N. Phan, Lillie S. Wendt, Charlotte A. Rose, Abigail Ali, David F. Carpio, Christine Cole, Paige Embley, Jack E. Hoskins-Harris, Delia Johnson, Noelle Ledoux, Hannah W. Lwin, Sarah Salah, Erin Weisbart, Stacey J. Criswell, Omar A. Quintero-Carmona

## Abstract

Myosin-X (MYO10) is an actin-based motor protein involved in cytoskeletal dynamics, membrane interactions, and integrin-mediated adhesion. To investigate MYO10’s cellular roles, we generated MYO10 knockdown (MYO10^KD^) HeLa and COS7 cell lines using lentiviral shRNA. Compared to wild-type cells, both MYO10^KD^ lines showed reduced proliferation and impaired cell migration in wound assays. There were fewer edge filopodia in HeLa cells. Additionally, MYO10^KD^ cells demonstrated increased spreading on laminin-coated substrates, suggesting altered integrin activation and cytoskeletal linkage. Our results reinforce MYO10’s importance in cell proliferation, adhesion, and migration; these MYO10^KD^ lines provide an accessible cell culture model for further study of MYO10.

---

In eukaryotic cells, both cell migration and the organization of internal components rely on a cytoskeletal system composed of filaments, motor proteins, and their associated regulators (reviewed in Machesky and Schliwa, 2000). The class X myosin proteins (MYO10) play roles in both cytoskeletal dynamics and membrane interactions (Kerber and Cheney, 2011; Tokuo, 2020). MYO10 facilitates the formation of filopodia (Berg and Cheney, 2002; Bohil et al., 2006; Tokuo et al., 2007), promotes directed migration (Arjonen et al., 2011; Hwang et al., 2009), and links actin filaments to membrane-bound receptors such as integrins (Cox et al., 2002; He et al., 2017; Toyoshima and Nishida, 2007; Zhang et al., 2004). Given these roles, MYO10 is thought to contribute to spatially coordinated signaling events essential for cell proliferation, adhesion, and motility. To better understand the mechanistic contributions of MYO10 in these contexts, we generated and characterized MYO10 knockdown cell lines using lentiviral delivery of short hairpin RNA (shRNA) resulting in genomic integration (Dull et al., 1998). By doing so, we aimed to generate an accessible cell culture model system for investigating the roles of MYO10 in cytoskeletal organization, migratory capacity, and adhesive behavior. These cell lines also provide a useful platform for studying the functionality of specific MYO10 domains and sequences through ectopic expression of fluorescently tagged or mutant constructs.

Cell culture model systems are more useful for studying the behavior of fluorescently tagged proteins-of-interest if the endogenous expression of that protein is reduced. By minimizing the confounding behavior of the native protein, observed phenotypes can be attributed to the exogenously expressed protein. Although embryonic fibroblasts isolated from MYO10 knockout mice (MYO10^KO^ MEFs) are available (Heimsath et al., 2017; Yim et al., 2023), we were unable to achieve robust transient transfection in MYO10^KO^ MEFs using the lipid-based methodology available to us. Commonly available mammalian cell culture models such as HeLa cells (Scherer et al., 1953), a human cervical carcinoma line, and COS7 cells (Gluzman, 1981), an African green monkey kidney epithelia line, are easily transfectable and have been used extensively to study MYO10 function (Berg and Cheney, 2002; Bishai et al., 2013; Kerber et al., 2009; Raines et al., 2012; Tokuo and Ikebe, 2004). These cell lines are also amenable to lentiviral delivery of shRNA-containing plasmids, which result in low-frequency genomic incorporation of lentiviral DNA. If the delivered plasmid also contains a puromycin mammalian antibiotic resistance marker, then semi-clonal lines can be identified when cultured in the presence of the antibiotic. Antibiotic-resistant sub-cultures can then be screened for decreased MYO10 mRNA and protein expression (Moffat et al., 2006). Verification of knockdown was done via qPCR analysis of MYO10 mRNA levels (Figure 1A) and quantitative western blot analysis of MYO10 protein levels (Figure 1B). Expression was reduced by >90% in shRNA-expressing HeLa and COS7 cell lines (MYO10^KD^) compared to their wild-type parental lines (Student’s t-test, p<0.05).

**Figure 1:**
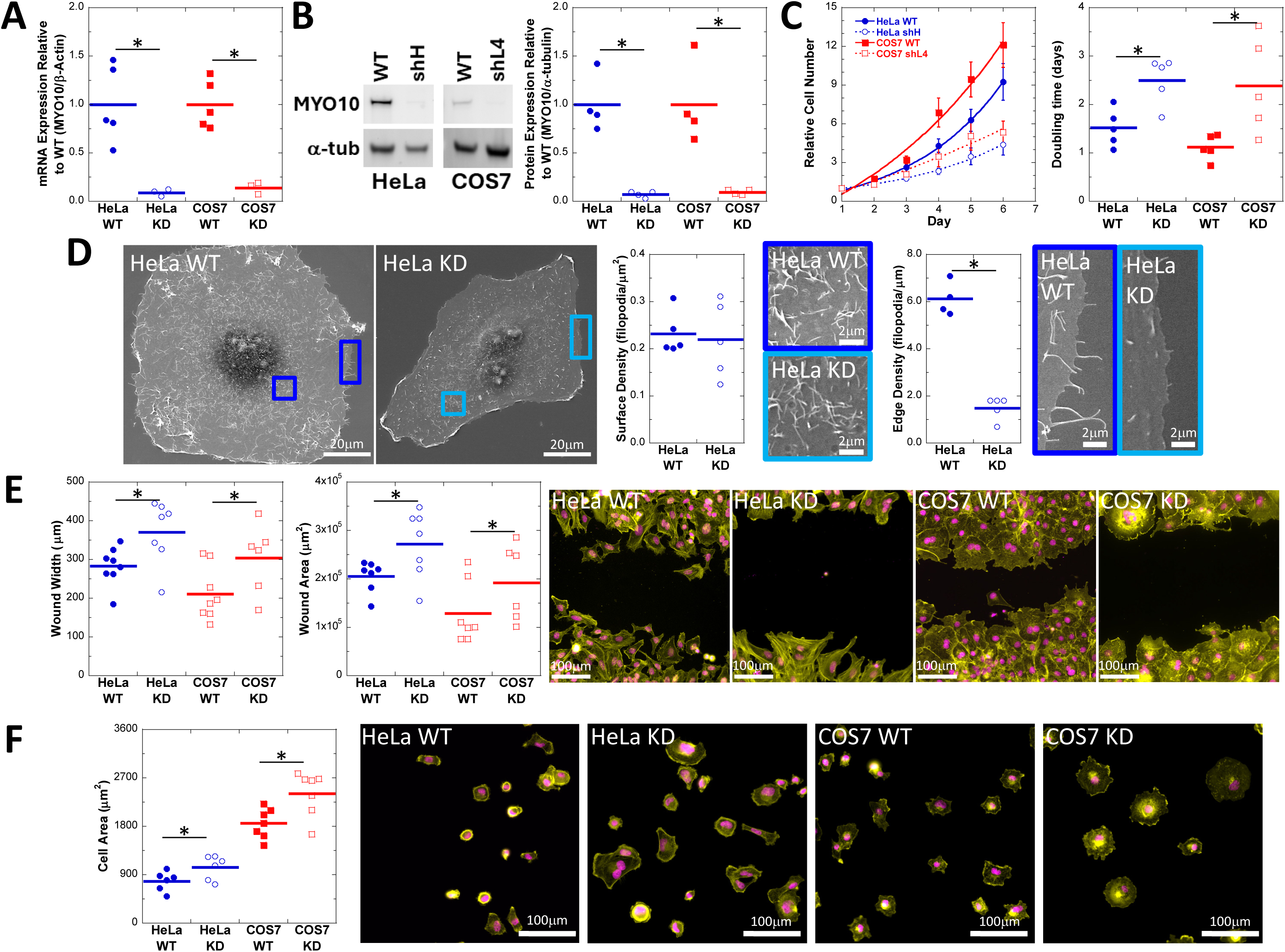
Decreased MYO10 expression in two different cell lines stably expressing shRNA results in proliferation, motility, and attachment defects compared to wild-type parental cell lines. Semi-clonal HeLa and COS7 lines isolated by puromycin selection show decreased MYO10 mRNA and protein expression as measured by **(A)** qPCR or **(B)** quantitative western blotting. The MYO10 band is nearly undetectable in a representative western blot. Blots against α-tubulin were used as a standard for quantification. **(C)** Both HeLa and COS7 MYO10^KD^ cells proliferate more slowly than their parental lines. The curve is a fit to a single exponential growth, and slower proliferation was quantified as the increased doubling-time for the shRNA expressing cells. **(D)** MYO10^KD^ HeLa cells maintain the ability to form dorsal surface filopodia with no change in filopodial density, but do show decreased edge filopodia density. The colored boxes on the lower magnification images correspond to the borders of the higher magnification images. **(E)** Cell migration following wounding is impaired in MYO10^KD^ cells, as demonstrated decreased wound width or wound area 18h after monolayer wounding. **(F)** Cell spreading on laminin-coated surfaces is enhanced in MYO10^KD^ cell lines, as illustrated increased cell area 2 hours after initial plating. *Quantitative and statistical representations:* For all plots, circle and square markers represent biological replicates, horizontal bars represent the mean across replicates, and asterisks represent Student’s t-test with p<0.05. For the growth curve plot in (C), markers represent the average of biological replicates and traces represent fitting the averaged data to single exponential growth. Representative images in (E) and (F) are stained for actin (ALEXA488-phalloidin, yellow) and DNA (DAPI, magenta).

We assayed the HeLa and COS7 MYO10^KD^ cell lines for their growth characteristics (Figure 1C). When doubling time was calculated from fitting growth data to single exponential growth, the observed doubling time was significantly slower by >1 day in each MYO10^KD^ cell line compared to its MYO10^WT^ counterpart (Student’s t-test, p<0.05). MYO10 is required for proper centromere and spindle orientation through its interaction with microtubules through the MyTH4 domain (Kwon et al., 2015; Toyoshima and Nishida, 2007; Weber et al., 2004; Woolner et al., 2008). Without proper spindle orientation, symmetrical chromosome segregation is impaired, and proliferation could be reduced. This slowed growth pattern of MYO10^KD^ cells is consistent with other findings where expression changes in MYO10 led to changes in cell proliferation (Mayca Pozo et al., 2023; Mayca Pozo et al., 2021).

HeLa cells generate filopodia on their dorsal surfaces and along their edges, while COS7 cells do not generate dorsal filopodia and have very few edge filopodia (Berg and Cheney, 2002; Bohil et al., 2006). Previous reports described differences in filopodia formation following transient small interfering RNA treatment to reduced MYO10 expression (Bohil et al., 2006). We used scanning electron microscopy to assay HeLa MYO10^KD^ cells for their ability to generate dorsal and edge filopodia (Figure 1D). We observed a significant decrease in the density of filopodia along the periphery of HeLa MYO10^KD^ when compared to HeLa MYO10^WT^ cells. However, analysis of dorsal surfaces showed no difference in filopodial densities between HeLa MYO10^KD^ and HeLa MYO10^WT^ cells, which differs from previous reports (Bohil et al., 2006; Pi et al., 2007). It is possible that the acute loss of MYO10 by transient siRNA transfection leads to different behavior than cells stably expressing shRNA, as cells stably expressing shRNA may have upregulated compensatory pathways that lead to dorsal filopodia formation (El-Brolosy and Stainier, 2017). It is also possible that the HeLa cell lines used in this study are potentially quite different from the HeLa cells used previously. There is a growing literature demonstrating that different HeLa cell isolates have different genomic signatures which could result in differences in phenotypes and behaviors (Frattini et al., 2015; Hu et al., 2019; Landry et al., 2013; Liu et al., 2019).

We used a wound-healing assay (Liang et al., 2007) to examine the effect of MYO10 knockdown on collective cell migration. Briefly, cells were grown to near-confluency on coverslips, the monolayer was disrupted by gentle scratching with a pipet tip, and the wounds were fixed and examined following 18 hours of recovery. Recovery was measured by either quantifying the width of the wound, or by assaying the area of the wound visible in each image (Figure 1D). Both MYO10^KD^ cell lines migrated into the wound more slowly than their MYO10^WT^ counterparts as illustrated by wider wound widths and by larger wound areas (p < 0.05). Our results are consistent with previous findings showing a reduction in cell motility following MYO10 knockdown as represented by decreases in cell migration across multiple cell types with reduced MYO10 expression (Makowska et al., 2015; Tokuo et al., 2018; Yu et al., 2015). These differences in behavior could be due to changes in filopodial assembly dynamics or to differences in the cells’ ability to adhere to substrates due to decreased MYO10 function. MYO10’s role in interacting with the extracellular matrix to form adhesions and influence motility makes it an integral component of the cell’s migratory machinery (He et al., 2017; Yu et al., 2015; Zhang et al., 2004). Knocking down MYO10 greatly reduces the interactions between cell and matrix, potentially making it more difficult for cells to move.

MYO10 serves as a link between the actin cytoskeleton and extracellular matrix proteins through integrin-mediated adhesions (Alieva et al., 2019; Zhang et al., 2004). Laminin is one such extracellular matrix protein. We used cell spreading assays on laminin-coated surfaces to evaluate the impact of decreased MYO10 expression on integrin-mediated attachment for both HeLa and COS7 cells. MYO10^KD^ and MYO10^WT^ cells were allowed to adhere to laminin-coated coverslips for 2 hours prior to fixation and phalloidin staining. The cells were then imaged and their attachment area was quantified (Figure 1F) using Ilastik (Berg et al., 2019) to identify cell area. We observed a significant increase in cell spreading for either MYO10^KD^ line compared to its wild-type parental line. Previous studies have also reported similar increases in cell spreading with decreased MYO10 expression (Bohil et al., 2006). Integrin activation and adhesion may be functioning as a governor, slowing the speed of cellular protrusion through the connection to the cytoskeleton via MYO10 (Courson and Cheney, 2015; Watanabe et al., 2010). MYO10 knockdown has been shown to impair integrin activation at filopodia tips, even though integrins remain localized along the filopodia shaft (Miihkinen et al., 2021). This impaired spatial regulation of integrin activity and decreased linkage between the integrins and the cytoskeleton may disrupt normal responses to the extracellular matrix, contributing to the enhanced spreading phenotype observed here. Our findings are consistent with the two-step model proposed by Miihkinen et al., where MYO10 tethers integrins at filopodia tips to facilitate talin-mediated activation (Miihkinen et al., 2021). MYO10 depletion would thus dysregulate this process, altering adhesion dynamics and spreading behavior (Yu et al., 2015).

Taken together, the phenotypes of these HeLa and COS7 MYO10^KD^ cells align with other previous reports of MYO10 knockdown (Bohil et al., 2006; Makowska et al., 2015), and knockout cell phenotypes (Heimsath et al., 2017; Ou et al., 2022; Tokuo et al., 2018). These new cell lines will aid in the study of MYO10 as they can be easily transfected with fluorescently tagged and modified versions of MYO10, allowing any scientist with cell culture and fluorescence imaging capabilities to investigate how particular changes to wild-type MYO10 structure influences MYO10. As lentiviral shRNA delivery systems are available for much of the mouse and human genome (Moffat et al., 2006), generation and characterization of knockdown cell lines could provide other undergraduate scientists with an opportunity for authentic contribution to broader scientific understanding for other genes, just as this project has for University of Richmond students gaining experience in quantitative imaging approaches for studying cell function.

## Materials and Methods

### COS7 and HeLa cell culture

Cos7 (Gluzman, 1981) and HeLa (Scherer et al., 1953) cells were cultured in growth media: DMEM high glucose (Gibco) supplemented with 10% Fetal Bovine Serum (Benchmark FBS, GeminiBio), and antibiotics (50U/ml penicillin, 50μg/ml streptomycin, Gibco). Cells were maintained in a humidified incubator at 37°C with 5% CO_2_ concentration and passaged using 0.25% trypsin-EDTA (Gibco).

#### Lentivirus and stable cell line production

HEK293FT cells (a gift from Benjamin Major, UNC Chapel Hill) were grown in growth media. The cells were maintained at 37°C in a humidified environment supplemented with 5% CO₂. To generate virus, Lipofectamine 3000 (Invitrogen) was used as the transfection reagent, and cells grown in T75 plates pCMV-VSV-G (Stewart et al., 2003), pRSV-Rev, pMDLg/pRRE (Dull et al., 1998) (Addgene plasmid # 8454, 12253, and 12251, respectively), pLKO.1 plasmid TRCN0000123087 (Moffat et al., 2006) targeting the human MYO10 sequence (Sigma), and a small amount of pEGFP-C1 to verify transfection. Twenty four hours later, media containing the virus was collected and replaced with fresh media. Cellular debris was removed from the virus-containing media by centrifugation at 2000 x g for 5 minutes at 4°C prior to infection.

HeLa cells and COS7 cells were plated in T25 flasks and grown to confluency. They were then incubated with lentivirus media diluted into DMEM growth at a 1:4 or a 4:1 viral media:growth media ratio. The media was supplemented with 8μg/ml hexadimethrine bromide and the infected plates were incubated overnight. The following day, the cells in the T25 were transferred to 150mm cell culture dishes and grown in media containing 2µg/ml puromycin. Media was replaced every two days to remove debris for dying cells. The plates were examined for clusters of isolated cells assumed to be colonies of cells where the DNA containing the shRNA cassette and the puromycin resistance gene had been incorporated into the cells’ genome. Eight to ten days after infection, clusters were isolated by placing a cloning ring in sterile petroleum jelly surrounding a colony. The colony was trypsinized, transferred to one well of a 6-well dish, and cultured in puromycin-containing growth media. The cultures were grown and expanded for 10 more days prior to verification of MYO10 knockdown by qPCR and western blotting. At this point, validated cultures of HeLa cells and COS7 cells were designated “passage #1,” and aliquots were frozen in DMEM growth media supplemented with 10% cell culture-grade DMSO (Sigma) and 40% fetal bovine serum.

#### Quantitative Real Time PCR sample preparation and analysis

RNA was isolated from T25-sized cultures of HeLa or COS7 cells using the RNeasy Plus kit (Qiagen) according the supplied protocol for purification of RNA from animal cells. In addition to the standard protocol, we also treated the cells to a Qiashredder step before loading the cell lysate onto the purification column, and we treated the lysate with an on-column DNAse digestion prior to washing and eluting the RNA. RNA concentration was estimated using a Nanodrop spectrophotometer, aliquoted, and frozen at -80°C.

cDNA was synthesized using the M-MLV reverse transcriptase kit with random hexamer primers, following the standard protocol, with a starting amount of 750µg RNA (Invitrogen). qPCR samples were prepared using the Luna Universal qPCR Master Mix (New England Biolabs) and run on a Bio-Rad CFX96 real time PCR machine, and assayed for MYO10 expression and β-actin expression. Standard curves for both MYO10 and β-actin were generated and run in parallel with the cell line cDNA samples—all samples in triplicate. MYO10 expression relative to β-actin expression was quantified using Bio-Rad CFX Manager software and is expressed relative to the wild-type cell MYO10/β-actin ratio for the relevant knockdown line.

#### Western blotting

Cells were lysed in sample buffer containing protease inhibitors (50 mM Tris HCl, pH 7.4, 0.5 M NaCl, 0.2% SDS, 1 mM EDTA, 1 mM DTT, 10 μg/ml aprotinin, 10 μg/ml leupeptin, 1 mM PMSF) at a concentration of 2×10^6^ cells/ml. After adding an appropriate volume of 5x Lamelli sample buffer, the samples were heated to 95°C for 5 minutes, and then allowed to cool to room temperature. Samples were loaded onto a 4-12% Bis-Tris NuPAGE gel, and run at ∼180V until the dye front reached the bottom of the gel. Proteins were transferred to Immuno-Blot Low Fluorescence PVDF Membrane, using an XCell II blotting apparatus (Invitrogen) in NuPAGE transfer buffer with 20% methanol for 60 minutes at 30V. Transfer was verified by Ponceau staining (0.5% Ponceau Red-S in 2% Acetic acid). Membranes were incubated in blocking buffer (1% Casein in TBS) for 1 hour prior to incubation with rabbit α-MYO10 antibody (Abcam #ab224120, 1µg/mL in blocking buffer) overnight at 4°C with gentle agitation. The blots were then washed 4 times in TBST for 10 minutes, and then incubated with hFAB™ Rhodamine α-tubulin (part#12004166 and 1:5000) an donkey α-rabbit ALEXA647 antibody (Jackson Immunoresearch, 0.15µg/mL in blocking buffer) for 1 hour. Blots were washed 4 times in TBST for 10 minutes each prior to visualization using a BioRad Chemidoc MP system using the CY5 and Rhodamine filter sets to visualize MYO10 and tubulin staining, respectively.

#### Western blotting quantification and analysis

Band intensities were quantified using the “Analyze→Gels” tool in FIJI/ImageJ (Schindelin et al., 2012). The MYO10/tubulin ratio was calculated for each sample, where the average MYO10/tubulin ratio for the wild-type cells was set to a value of 1, and all other MYO10/tubulin ratios were standardized relative to this value.

#### Growth curve sample preparation, data acquisition, and analysis

The cell proliferation assay protocol was adapted from (Rago et al., 1990). Cells were plated in six 96-well cell culture plates at a density of 1000 cells/well in 100 μL of growth media and incubated in a humidified environment at 37°C with 5% CO₂. Empty wells were filled with 100 μL of sterile water to prevent media evaporation. Sufficient plates were set up at the beginning of the experiment so that one plate could be collected each day of the experiment. Each day, one plate was removed from the incubator, and media was discarded by gently shaking the plate over a sink, followed by blotting on a paper towel. The plate was then stored at -80°C. This process was repeated at the same time for subsequent days. All plates were kept at -80°C for an additional 24 hours following the final time day. Plates were thawed at room temperature for approximately an hour. After thawing, 100 μL of sterile water was added to each well. To induce lysis, plates were incubated at 37°C for one hour before being returned to -80°C overnight. Plates were removed from -80°C, thawed at room temperature for an hour, and stained with 100 μL of staining buffer (10mM Tris, 2M NaCl, 1mM EDTA, 2mM NaN_3_, 2.5μg/mL Hoescht 33342, pH 7.4). Plates were incubated at room temperature in the dark for 30 minutes before fluorescence measurement. Fluorescence intensity was recorded using a Spectramax M3 plate reader (Molecular Devices) with the following settings: excitation/emission range of 360-460 nm, no shaking, and a top-down detection mode (plate lids removed prior to reading). Data was saved and exported as Excel files for further analysis.

Raw absorbance data was pasted into a spreadsheet for analysis. Background absorbance values were determined using sterile water wells and subtracted from all sample wells. To calculate fold-growth, absorbance values for each condition were normalized to the average absorbance on Day 1. Average fold growth values across multiple experiments were plotted using KaleidaGraph software, and were then fit to a single exponential growth model. The exponential growth curve was used to determine the doubling time at Day 4.

#### SEM sample preparation, imaging, and filopodial density quantification

Before processing, a 6-well dish was prepared, where wild-type or MYO10 knockdown HeLa cells were plated on acid-washed(Berg et al., 2019), 18 mm round #1.5 coverslips at a concentration of 25,000 cells/well in growth media, and allowed to adhere overnight. The following day, media was removed, and samples were fixed in warm PBS containing 4% paraformaldehyde and 2% glutaraldehyde for at least 30 minutes with gentle rocking. Fixative was removed with three, 10-minute PBS washes. Samples were dehydrated via a series of ethanol washes at 30%, 50%, 70%, 95%, and 100% ethanol for 10 mins at each concentration. Samples were submerged in 100% ethanol and critical-point dried using a Samdri-795. After drying, samples were mounted on 12.2 mm diameter, 10 mm high aluminum mounts and sputter-coated with a Leica EM ACE600 ∼10nm of 60/40 gold/palladium to prevent the buildup of surface charges. Samples were then imaged using a JEOL JSM-IT700HR scanning electron microscope set to a working distance of 15mm, a standard point current (STD-PC) of 50, and a field of view of 128µmx96µm.

To reduce the potential for bias, SEM images were randomized using a “Blind Experiment” plugin in FIJI-ImageJ. Randomized images were then opened in FIJI. To calculate dorsal filopodia density, a region of interest (ROI) was highlighted using the “Rectangle” tool for each cell present in the image. The ROI was then enlarged, and dorsal filopodia were manually counted using the “Cell Counter” plugin. Filopodial surface density (filopodia/mm^2^) was then calculated through the counted filopodia by the area of the region of interest. To calculate edge density, the length of free cell edge was measured and the number of filopodia along that edge were counted (filopodia/μm).

#### Wound healing assay sample preparation & imaging

On day 1, both HeLa and COS7 wild-type and MYO10 knockdown cells were plated on square glass coverslips at a concentration of 120,000 cells per well in growth media and allowed to adhere overnight. On day 2, once confluency was verified, cells were scratched. Using a P10 pipette tip, the coverslip upon which the cells were growing on was scratched 5 total times. 3 of the scratches were done in parallel vertically, and 2 of the scratches were done in parallel horizontally. This allowed up 5 possible “lanes” of observation when image capturing. After scratching, the media from these wells was removed, and 3 mLs of Low-Serum Media was added to prevent cellular division. Wounds were allowed to heal for 18 hours, after which cells were then permeabilized in PBS with 0.5% Triton X100 for 5 minutes. Samples were stained for F-actin and DNA with 6.6nM ALEXA488-phalloidin and 15nM DAPI for 30 minutes. After four 5-minute PBS washes, the cells were mounted onto slides using Profade Glass. Images were captured using an Olympus UPlanSApo20x/0.75NA objective mounted on either an IX-71 or IX-83 stand, an EXFO mixed gas light source, a Chroma Sedat Quad filter set, and a Hamamatsu ORCA Flash 4v2 camera. The microscopy hardware and image acquisition settings were controlled by Metamorph software. Typical exposure times were under 30ms for DAPI and under 150ms for ALEXA488-phalloidin.

#### Wound healing assay quantification, widths

Using the straight line tool in FIJI/ImageJ, ten lines were traced from one side of the wound to the other, approximately perpendicular to the wound edge for each image. Line lengths were recorded in pixels, converted to microns, and averaged for each day. A minimum of 10 images were captured for each condition on each day.

#### Wound healing assay quantification, area

Wound area was completed using both Ilastik (Berg et al., 2019) and FIJI/ImageJ. With Ilastik, the images captured on the channel visualizing F-actin were uploaded to the application. “Cell” and “background” pixels were identified using the “Pixel Classification” segmentation workflow, with all options selected in the “Feature Selection” step. The classifier was trained using 15-20% of the total number of pictures in a set. Scratches from each day, cell type, and condition were trained individually. Simple segmentations were exported as TIF files. The resulting segmented files were thresholded in FIJI/ImageJ, the wound areas were measured for each image, and averaged together for each condition on each day. A minimum of 10 images were captured for each condition on each day.

#### Spreading assay sample preparation & imaging

Prior to plating, acid-washed, 22mm square #1.5 coverslips were placed in a 6 well dish, coated with 10µg/mL laminin in PBS for 30 minutes, and then washed 3x with PBS. Cells were lifted using 0.25% trypsin with EDTA, and then resuspended at a concentration of 20,000 cells/mL in DMEM supplemented with 10% fetal bovine serum, penicillin, and streptomycin. 3 mL of the cell mixture were added to each well for a total of 60,000 cells/well, and allowed to adhere for the specified amount of time. Once the time-course ended, the media was removed and the cells were fixed in PBS with 4% paraformaldehyde for 20 minutes. Cells were then permeabilized in PBS with 0.5% Triton X100 for 5 minutes. Samples were stained for F-actin and DNA with 6.6nM ALEXA568-phalloidin and 15nM DAPI for 30 minutes. After four 5-minute PBS washes, the cells were mounted onto slides using Profade Glass. Images were captured using an Olympus UPlanSApo20x/0.75NA objective mounted on either an IX-71 or IX-83 stand, an EXFO mixed gas light source, a Chroma Sedat Quad filter set, and a Hamamatsu ORCA Flash 4v2 camera. The microscopy hardware and image acquisition settings were controlled by Metamorph software. Typical exposure times were under 30ms for DAPI and under 150ms for ALEXA568-phalloidin.

#### Spreading assay quantification, area

Measurement of cell area was completed using both Ilastik and FIJI/ImageJ. With Ilastik, the images captured on the channel visualizing F-actin were uploaded to the application. “Cell” and “background” pixels were identified using the “Pixel Classification” segmentation workflow, with all options selected in the “Feature Selection” step. The classifier was trained using 15-20% of the total number of pictures in a set. Samples from each day, cell type, and condition were trained individually. Simple segmentations were exported as TIF files. The resulting segmented files were thresholded in FIJI/ImageJ, cell areas were measured for each image, and averaged together for each condition on each day. A minimum of 10 images were captured for each condition on each day.

#### Replication and statistical analysis

For qPCR, each marker represents the average of 2-3 technical replicates for a particular experiment (n = 3 or 4 experiments). For western blotting, each marker represents a single sample for a particular experiment (n =4 experiments). For growth curves and doubling time calculations, each marker represents the average of 5-7 different experiments where each experiment contained triplicate technical replicates. For dorsal filopodia density, each marker represents the average of >15 measured cells (n = 5 experiments). For wound width, each marker represents the average of >50 width measurements across multiple wounds (n = 6, 7, or 8 experiments). For wound area, each marker represents the average of >10 different images of wounds (n = 6 or 7 experiments). For spreading area, each marker represents the average of >90 cells (n = 6 or 7 experiments). Statistical comparisons between conditions within a cell type (wild-type versus knockdown) were carried out using a Student’s t-test.

## Acknowledgements

The results in this report were generated primarily by undergraduate students working in the Quintero lab and the Spring BIOL317 Mechanochemical Cell Biology Course at the University of Richmond. OAQ-C generated the lentivirus and established the cell lines, CEC validated the cell lines using qPCR and western blotting, JAM completed the growth assays, VMNP completed the scanning electron microscopy analysis, and CEC & JAM collaborated on the scratch analysis. Once the cell lines were established the students in BIOL317 used the lines as part of a course-based undergraduate research experience where they became proficient at collecting fluorescence microscopy data, and basic segmentation and quantitative image analysis under the guidance of SJC, EW, and OAQ-C. JAM was the BIOL317 teaching assistant responsible for cell line maintenance and sample preparation. The students chose to analyze cell spreading and other measures of size & shape. Validation of the results came from multiple teams of students using different instruments to collect and analyze images from the same sets of slides. The students then compared results across all teams within the class. To get exposure and immersion to the cell science field, the students presented their results in a “Zoom lab meeting” at the end of the semester to Hijab Fatima (Columbia University), Jordan Beach (Loyola University Chicago), Richard Cheney (UNC Chapel Hill), Tom Pollard (Yale University), Derek Applewhite (Reed College), Meg Titus (University of Minnesota), Kendall Stewart (UC Davis), Gabe Offenback (UNC Chapel Hill), Graham Johnson (Allen Institute for Cell Science), Uri Manor (UC San Diego), Sarah Heisler (The Ohio State University), Adelaide Masterson (Latham BioPharm Group), Anthony Isenhour (Yale University), Dan Kiehart (Duke University), and John Peters (Harvard University). The BIOL317 team would like to thank these scientists for their time, their support, and their insight. Their participation was crucial to giving the students a sense that their efforts have value to the greater cell science community. We thank the University of Richmond School of Arts & Sciences for establishing and maintaining the <u>Biological Imaging Lab</u>. Laboratory courses based in modern microscopy approaches are uncommon at the undergraduate level and prepare Richmond students for future success in science.

## Author Contributions

Joanna A. Mas: data curation, formal analysis, investigation, methodology, supervision, validation, visualization, writing-original draft, writing-review & editing

Chase E. Cristella: data curation, formal analysis, investigation, methodology, validation, visualization, writing-original draft, writing-review & editing

Vu M.N. Phan: data curation, formal analysis, investigation, methodology, validation, visualization, writing-original draft, writing-review & editing

Lillie S. Wendt: data curation, formal analysis, investigation, methodology, visualization, writing-original draft, writing-review & editing

Charlotte Rose: data curation, formal analysis, investigation, methodology, visualization, writing-original draft, writing-review & editing

Abigail Ali: data curation, formal analysis, investigation, methodology, visualization

David F. Carpio: data curation, formal analysis, investigation, methodology, visualization

Christine Cole: data curation, formal analysis, investigation, methodology, visualization

Paige Embley: data curation, formal analysis, investigation, methodology, visualization

Jack Hoskins-Harris: data curation, formal analysis, investigation, methodology, visualization

Delia Johnson: data curation, formal analysis, investigation, methodology, visualization

Noelle Ledoux: data curation, formal analysis, investigation, methodology, visualization

Hannah Lwin: data curation, formal analysis, investigation, methodology, visualization

Sarah Salah: data curation, formal analysis, investigation, methodology, visualization

Erin Weisbart: formal analysis, methodology, supervision, writing-review & editing

Stacey J. Criswell: conceptualization, funding acquisition, methodology, project administration, resources, supervision, writing-review & editing

Omar A. Quintero-Carmona: Conceptualization, data curation, formal analysis, funding acquisition, investigation, methodology, project administration, resources, supervision, validation, visualization, writing-original draft, writing-review & editing

